# Hyperspectral Imaging to Quantify Nodules and Detect Biological Nitrogen Fixation in Legumes

**DOI:** 10.1101/2025.07.25.666867

**Authors:** Yuchen Wang, Kithmee De Silva, Di Song, Mohammed Kamruzzaman, Matthew D. Brooks

**Author notes:** These authors contributed equally to this work (Yuchen Wang, Kithmee De Silva, Di Song).

## Abstract

Legume root nodules are important for biological nitrogen fixation, a process critical for plants to gain additional nitrogen from the environment. Nodule quantification is valuable for evaluating nitrogen fixation efficiency, assessing symbiotic relationships, monitoring responses to nitrogen, and supporting genetic studies on legume adaptation and productivity. However, accurate quantification of root nodules is difficult and time-consuming due to the complexity of the root system and soil interference. Here, we explore the utility of hyperspectral imaging as a non-destructive tool to detect active fixing root nodules with minimal preparation and show that we can differentiate nodules and root tissues through unique spectral signatures while also distinguishing between fixing and non-fixing nodules. We applied deep learning techniques to develop an automated nodule counting pipeline adaptable across different legume species and under diverse growth conditions. This approach eliminates the need for labor-intensive counting and enables the detection of nodules embedded within dense root tangles with high accuracy. This automated hyperspectral approach offers a promising alternative to support assessments of nodule abundance and their activity across legume species grown under various environments.

## Introduction

Legumes (Fabaceae or Leguminosae family), the third largest family of angiosperms, are known for their worldwide distribution and agricultural value. Their ability to form a symbiotic relationship in root nodules with nitrogen-fixing rhizobia in the soil distinguishes them from non-nodulating plants. This association enables the conversion of atmospheric nitrogen (N_2_) into ammonia (NH_3_), a biologically usable form of nitrogen that supports plant growth. In return, the plant supplies the bacteria with carbon derived from photosynthesis. This process, known as biological nitrogen fixation (BNF), reduces reliance on synthetic nitrogen fertilizers and allows legumes to thrive in nitrogen-poor soils, contributing to sustainable agriculture (Peoples et al., 1995). Studies suggest that BNF may give legumes a competitive advantage over non-fixing plants under future elevated CO_2_ levels as the additional nitrogen gained helps maintain the plants’ carbon-to-nitrogen balance (Rogers et al., 2009).

This symbiosis is facilitated by specialized root structures called nodules, which house the bacteria and create a microenvironment suitable for the oxygen-sensitive enzyme nitrogenase. Leghaemoglobin, a heme protein found in nodules, binds free oxygen to maintain the low-oxygen conditions required for nitrogenase function while transporting O^2^ to bacteria for respiration (Appleby, 1984; Ott et al., 2005). However, nodulation and nitrogen fixation are known to vary, both within and among legume species (Bourion et al., 2007; Hardarson et al., 1993; Iqbal et al., 2022). This variability results from a complex interplay of genetic and environmental factors, as the host primarily shapes nodule morphology and structure while the efficiency of nitrogen fixation is largely determined by the bacterial symbiont partners (Mortier et al., 2012; Sprent, 2007). Different legume species (and even different genotypes within a species) can exhibit distinct nodule characteristics, including variations in number, size, weight and morphology (Sprent et al., 2017). For instance, some legumes develop determinate nodules with a spherical, fixed size, while others form cylindrical indeterminate nodules that continue to grow and display meristematic activity. These inherent differences not only affect the total capacity for nitrogen fixation but also influence the efficiency and longevity of the symbiosis (Bourion et al., 2007; Tajima et al., 2007). Nodule phenotypic traits such as number, size, biomass, internal hue/pigmentation, and external morphology have been frequently used as proxies for BNF capacity (Hensley et al., 2021; Tajima et al., 2007). Traditional methods, such as manual counting and nodule area estimation (Dias et al., 2017; Tajima et al., 2007) have been complemented by advances in imaging that enable semi-automated nodule quantification through both destructive and non-destructive approaches. Destructive methods include direct quantification of isolated nodules (Barbedo, 2012; Dias et al., 2017; Vikman and Vessey, 1993) and microscopic assessment on nodule cross sections (Tajima et al., 2007; TSYGANOV et al., 2002), while non-destructive approaches involve imaging minimally disturbed field-grown roots using minirhizotron platforms (Rowland et al., 2015), digital scans of washed intact root systems (Lira and Smith, 2000), and rhizoboxes (Nagel et al., 2012). Despite these improvements in throughput, they remain time-consuming, labor-intensive, and are confounded by the high variability in nodule size, shape and texture, and the inherent clustered nature of nodules. Also, these morphological indicators often correlate poorly with actual nitrogen fixation, in part because nodules at different developmental stages, including senescent ones, can appear similar yet differ drastically in metabolic activity (Dupont et al., 2012; Fernández-Luqueño et al., 2008; Vikman and Vessey, 1993). To address this, biochemical assays such as leghaemoglobin extraction (Wilson and Reisenauer, 1963) and acetylene reduction (Hardy et al., 1968; Vance et al., 1979) assess nodule function more directly. However, these assays are destructive, involve hazardous chemicals, and lack precision across diverse nodule types and developmental stages. Collectively, these constraints highlight the need for non-destructive, functionally informative, and scalable approaches to quantify BNF accurately.

Unlike conventional RGB imaging, which relies on morphological cues, hyperspectral imaging (HSI) captures reflectance spectra across hundreds of narrow, contiguous wavelengths, enabling the detection of subtle biochemical and structural differences. This spectral sensitivity is particularly relevant given that internal pigmentation gradients in nodules, linked to nitrogen-fixing activity, may produce detectable signatures in hyperspectral data (Chung et al., 2020; Hensley et al., 2021). HSI has been effectively used to monitor plant physiological processes such as stress responses (Behmann et al., 2014) and disease progression (Berdugo et al., 2014), indicating its potential for assessing nodule function.

Complementary advances in deep learning, especially convolutional neural networks (CNNs), have enabled accurate nodule segmentation from high-resolution RGB images of soybean roots (Woo et al., 2023). Utilizing CNNs, the YOLO (You Only Look Once) series, a deep learning object detection algorithm, stands out for its speed and accuracy. Since its introduction as a single-stage detection algorithm, YOLO has rapidly evolved to include multi-scale feature extraction, attention mechanisms, and transformer-based modules (Redmon et al., 2015). YOLOv11 builds upon these advancements to achieve high precision with reduced computational load, making it well-suited for root nodule detection in complex soil environments. Therefore, integrating spectral information with deep learning-based models can improve both the precision and the physiological relevance of an automated nodule detection model.

In this study, we present an HSI-based pipeline integrated with deep learning to non-destructively identify and quantify root nodules. By leveraging the spectral sensitivity of HSI and the strong object detection capabilities of the deep learning algorithm YOLOv11, we developed a model capable of distinguishing actively nitrogen-fixing nodules under varied soil conditions. Then, we applied our detection pipeline to five legume species to demonstrate its broad applicability across a range of morphologies and nodule types. Finally, we evaluate model performance, explore spectral features associated with functional nodules, and demonstrate the scalability of this approach for high-throughput root phenotyping in a real research experiment.

## Materials and Methods

### Plant materials and growth conditions

Five legume species, common bean (*Phaseolus vulgaris)*, soybean (*Glycine max*), cowpea (*Vigna unguiculata*), *Medicago truncatula,* and red clover (*Trifolium repens*) were used in this study. Scarified and imbibed medicago seeds were vernalized for 48 h in the dark at 4 °C, incubated for 24 h at room temperature and germinated on 1% agar plates vertically in a growth chamber under 200 ümol m^-2^ s^-1^ for 72 h. The seedlings were transferred to 4” pots containing either sterile potting medium (BM6 All-Purpose, Berger, Québec, Canada) or 1:1 mixture of coarse calcined clay (Turface MVP, PROFILE, Buffalo Grove, IL, USA) and fine clay (Greens Grade, PROFILE, Buffalo Grove, IL, USA). Common bean seeds were surface sterilized and germinated on dampened filter paper placed on a petri dish for 48 h. The seeds with radicles were transplanted in 6” pots with 1:1 mixture of coarse calcined clay (Turface MVP, PROFILE, Buffalo Grove, IL, US) and fine silica sand. Soybean, cowpea and clover seeds were directly sown in 6” pots with mixture of coarse calcined clay and fine silica sand. *Medicago* and clover plants were grown under a light intensity of 300 ümol m^-2^ s^-1^ under 8/16 h day/night cycle, 24/22 °C day/night temperature regime and 65% humidity while common bean, soybean and cowpea were grown under 800 ümol m^-2^ s^-1^ under 12/12 h day/night cycle, 28/25 °C day/night temperature regime and 65% humidity. The plants were inoculated with rhizobacteria either sowing into the soil or by directly appling to the base of the plantlet 3 days after transplanting (details of rhizobial strains in Table S1). The plants were supplemented with 0.1x fertilizer (Schultz All Purpose Plant Food 20-20-20, Knox Fertilizer, Knox, IN, USA) once a week to promote nodule formation under nitrogen-limiting conditions. All plants were watered every two days to maintain uniform soil moisture.

### Rhizobox design

The rhizobox was assembled using a darkened acrylic back panel and a transparent front panel made of glass, secured within a wooden frame with drainage holes at the base. The rhizobox has dimensions of 27 cm × 23 cm × 4 cm (height × width × depth). Soybean seedlings germinated in soil for five days were transplanted into rhizoboxes filled with either potting medium (BM6 All-Purpose, Berger, Québec, Canada) or fine silica sand. The rhizoboxes were positioned at a 45° angle with the glass panel facing downward to encourage root growth along the transparent surface. The glass surface was covered with aluminum foil to minimize light exposure to the roots.

### Nodule sampling and hyperspectral image collection

Roots were harvested and thoroughly rinsed with distilled water to remove residual soil particles, taking care not to damage the nodules. The harvested root can be stored in −80 °C if images are not planned to be taken within 24 hrs. Before imaging, excess water was gently removed while ensuring the roots remained moist to prevent desiccation. The roots were then carefully spread out to ensure even distribution and uniform height, minimizing nodule overlap and facilitating accurate identification. For samples grown in rhizoboxes, imaging was conducted directly through the glass panel. Hyperspectral data were collected from each root sample for subsequent analysis. Following imaging, nodules were manually counted with direct observation from the root to compare with model performance. A subset of roots was stored at −80 °C for biochemical assays.

### Nodule absorbance scan and leghaemoglobin quantification assay

Frozen root samples from 6-week-old common beans grown under minimal nitrogen in a 1:1 mixture of coarse calcined clay and fine clay were carefully separated into ‘green’ nodules, ‘pink’ nodules and root tissue. For each type, 1 g of tissue was weighed and crushed in liquid nitrogen. Total protein was extracted using 1 mL of 1x phosphate buffered saline (PBS, Corning 21-040-CV). The samples were centrifuged at 10,000 g for 15 minutes to pellet the debris. The absorbance spectra of the clarified supernatant were measured from 400 to 1000 nm with 2 nm step increments using the TECAN Infinite 200 PRO series microplate reader.

The spectrophotometric leghaemoglobin assay was done as described by Senthilkumar et al. (Senthilkumar et al., 2021). The supernatant from each sample was diluted fourfold (1:4) with 1x PBS buffer. An equal volume of alkaline pyridine reagent (4.2 M Pyridine in 0.2 M NaOH) was added. This mixture was split into two aliquots, of which one was treated with a few crystals of sodium dithionate, and the other with potassium hexacyanoferrate. Absorbance was measured using the microplate reader at 556 nm and 539 nm for sodium dithionate-treated and potassium hexacyanoferrate-treated samples, respectively. Leghaemoglobin content was calculated by the following equation (where D is the dilution factor) and normalized for the initial tissue weight;

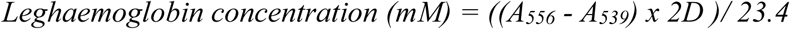

### Hyperspectral Camera Setup

Hyperspectral images were acquired using a line-scan benchtop HSI system (Resonon Inc., Bozeman, MT, USA) operated in a controlled dark environment. The system consisted of a 900-pixel line-scan hyperspectral camera (Pika L, Resonon Inc.) covering a spectral range of 374–1015 nm with a spectral resolution of 2.7 nm, yielding a total of 281 spectral bands. The imaging setup also included a linear translation stage for sample movement and a computer equipped with Spectronon Pro software (Resonon Inc.) for system control and data acquisition. The camera offered a maximum spatial resolution of 900 × 900 pixels and supported frame rates up to 249 fps. Prior to analysis, raw hyperspectral data were corrected for system noise and illumination variability by applying dark correction (removing sensor background signal) and white reference correction (normalizing reflectance using a calibrated white reference panel). These corrections were performed automatically using the software to ensure accurate reflectance spectra for all pixels across the image.

### Hyperspectral Imaging process

After acquiring full-range hyperspectral data, reflectance values at 660 nm, 550 nm, and 470 nm were extracted to generate standard RGB composite images. In parallel, reflectance at 515 nm, 560 nm, and 600 nm—wavelengths identified as showing spectral differentiation between root and nodule tissues—were selected to construct ‘enhanced images’. To enhance the contrast between nodules and root tissue, we computed a pixel-wise slope across these three hyperspectral bands. First, slope_1 was calculated between 515 nm and 560 nm. Next, slope_2 was calculated between 550 nm and 600 nm. The final pixel value for the enhanced image was obtained by subtracting slope_1 from slope_2. The resulting slope image was normalized to a grayscale range (0–255) to enhance nodule visibility and support segmentation.

### Model training and generation of validation data

After generating RGB and enhanced images, each image was manually annotated using the web-based annotation tool MakeSenseAI (Skalski, 2019). The total number of nodules identified per image was recorded to serve as a reference for model evaluation. The annotated label coordinates, along with the corresponding images, were used to train object detection models using YOLOv11. To account for physiological differences in nodule development, the dataset was divided into two categories: determinate and indeterminate nodules. Separate models were trained for RGB images and enhanced hyperspectral images within each nodule category to evaluate detection performance across both image types. A separate validation dataset that comprises RGB and enhanced images not seen during training was used to assess model performance (details of datasets in Table S2). For each validation image, the trained model outputs the total number of detected nodules, along with predicted bounding box coordinates and associated probability scores. A confidence threshold was applied to confirm valid nodule detections. To compare the predicted counts with image counts and visual counts, Pearson correlation analyses were performed using the cor() function in R (version 4.5.0). The relative accuracy of the predicted counts was calculated using the following equation:

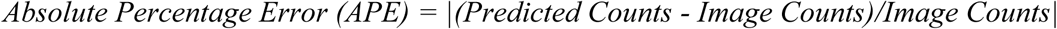

### Experiment Setup and Physiological measurements

Common bean and *Medicago truncatula* were grown in a factorial condition with varying nitrogen treatments and CO_2_ levels (details of the experiment setup in Table S3). After harvesting the roots, both shoots and root tissues were dried at 65 °C for 96 h for biomass quantification (root biomass include nodules). The nitrogen use efficiency (NUE) and nodule density were calculated using the following equations:

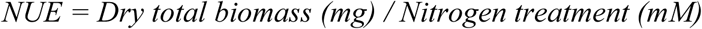

A three-way ANOVA followed by Tukey’s test was performed to assess the effects of nitrogen level, CO₂ level, and count type (predicted vs. visual) on nodule counts. Pearson correlation analyses were conducted using the cor() function in R (version 4.5.0) to evaluate the relationships between physiological measurements and nodule counts.

## Results

### Legumes exhibit species-specific nodule morphology

Five legume species were selected for this study, representing a range of nodule developmental types. These included three determinate species, common bean (*Phaseolus vulgaris*), cowpea (*Vigna unguiculata)* and soybean (*Glycine max*); two indeterminate species, *Medicago truncatula* and red clover (*Trifolium repens*). To capture a broader nodule physiological variation, experimental conditions including growing media, growth duration and nutrient supplementation regimes were varied while maintaining optimal growth conditions for each species (Table S1).

We observed noticeable differences in nodule size, density and morphology across the five legume species (Figure 1A), with a clear distinction between determinate and indeterminate nodule types (Figure 1B). Indeterminate nodules, as seen in *Medicago* and clover, formed elongated, branched nodules. Determinate nodules of common bean, cowpea and soybean were round and often formed clusters. Size of nodules also varied greatly within each plant individual due to their developmental stages. A consistent pattern across species was the external nodule coloration associated with nitrogen-fixing activity (Navascués et al., 2012). Pink coloration, typically associated with leghaemoglobin content, indicates active nitrogen fixation, while green or white nodules often indicate inactive or senescing nodules as the heme content degrades (Dupont et al., 2012). In all species except soybean, nodules typically showed pink and green hues externally. In contrast, soybean nodules appeared uniformly brown on the surface, but upon dissection, internal tissue color ranged from pink to green, reflecting different functional states.

**Figure 1.**
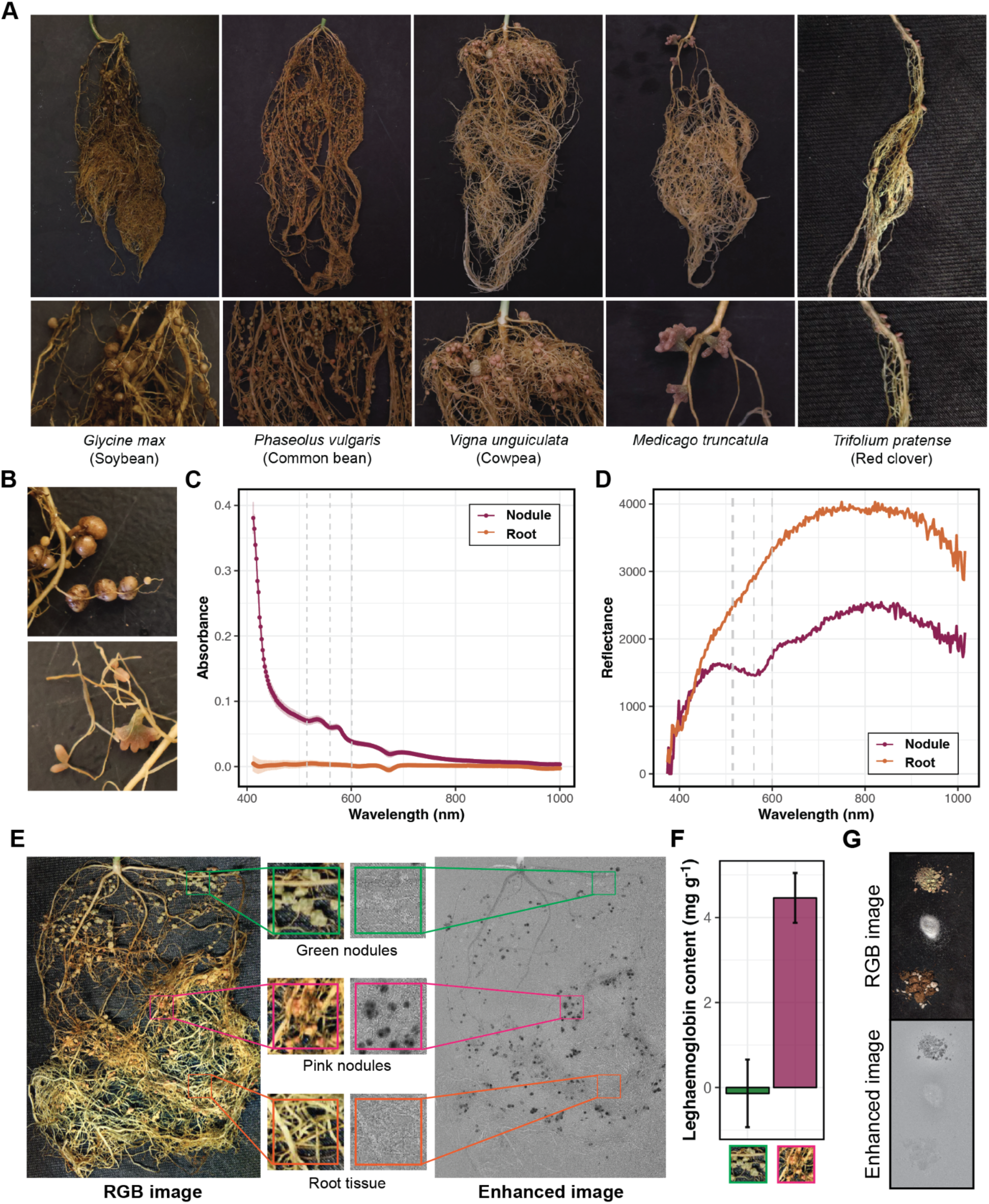
Hyperspectral imaging captures morphologically and physiologically diverse legume nodules through reflectance associated with leghaemoglobin. (A) Root systems and nodules of the five legume species selected for this study. (B) Representative images of indeterminate (top) and determinate (bottom) nodule types. (C) Absorbance spectra of protein extracts from isolated nodules and root tissue. (D) Reflectance spectra of different root tissue types. (E) RGB and enhanced images of a common bean root system, illustrating how different tissue types are represented in each image. (F) Quantification of legheamoglobin content in green and pink nodules in common bean (n=3). (G) Hyperspectral signatures of various potting media; clay, sand and potting mix (BM6), shown from top to bottom.

### Hyperspectral signatures capture biochemical differences between nodules and root tissue

Although legumes exhibit species-specific nodule morphology, nodules across species share a fundamental protein composition required to facilitate symbiosis and nitrogen fixation, and this protein profile is distinct from that of root tissues (Mergaert et al., 2020; Ott et al., 2005). As an initial assessment of whether biochemical and compositional differences between nodules and root tissue influence spectral properties, we separately isolated each tissue type and extracted total protein content. Absorbance scans of the protein extracts revealed distinctly different spectral profiles between the two tissues (Figure 1C), supporting the hypothesis that hyperspectral imaging may capture these differences in situ.

For imaging, we selected plants of each legume species aged 4-6 weeks with well-developed nodules and thoroughly cleaned the root systems to remove all potting media. First, to identify informative hyperspectral wavelengths, we performed full-spectrum reflectance scans using the Resonon HSI system. Regions of interest were visually selected based on RGB images to compare spectral reflectance across tissue types. This revealed distinct differential peaks at 515, 560, and 600 nm (Figure 1D). These three spectral bands were subsequently processed to generate an enhanced image (hereafter referred to as the “enhanced image”). These enhanced images helped the visual distinction of nodules from root tissues, where clusters of dark pixels (black dots) correspond to areas of active nodule presence, while root tissue blends into the lighter gray background. An example of an RGB image alongside its corresponding enhanced image after processing is shown in Figure 1E.

### Enhanced imaging selectively identifies functional/active nodules

An unexpected observation from enhanced images was the spectral difference between green and pink nodules on common beans, with green nodules appearing to be more similar to root tissue than nodules (Figure 1E). Given that green nodules are typically senesced (Navascués et al., 2012), the presence of active leghaemoglobin in pink nodules likely drives this spectral distinction, consistent with their characteristic coloration. In *Medicago* mutants such as *does not fix (dnf)*, which lack leghaemoglobin in their nodules (Starker et al., 2006) also showed no spectral differences compared to root tissue (<u>Figure S1</u>). Further quantification of leghaemoglobin content in nodules revealed significantly lower, almost undetectable levels of leghaemoglobin in green nodules (Figure 1F). The leghaemoglobin assay findings support that the spectral distinction is a result of this molecule, and also highlight a key feature of our approach, the selective identification of actively nitrogen-fixing nodules within the root system.

This approach effectively distinguished nodule tissue from surrounding root and background material with minimal cleaning of root systems for accurate imaging. Furthermore, the use of different growth media revealed that calcined clay emitted a reflectance similar to nodules (Figure 1G), making it challenging to work with unless thoroughly cleaned to prevent misidentification. However, general potting mix (BM6) and silica sand did not interfere with the nodule identification process (Figure 1G). Additionally, direct imaging of roots in rhizoboxes was feasible when using non-interfering substrates (<u>Figure S2</u>).

### Nodule-type specific training improves nodule detection

Given the clear separation of nodules in enhanced images, we deemed them to be well-suited for automated feature extraction, motivating us to apply deep learning–based models for nodule quantification. For this, following spectral enhancement to differentiate nodules from the background root tissue, images were preprocessed for object detection model development. Specifically, bands 450, 550, and 670 nm were combined to generate RGB images, while bands 515, 560, and 600 nm were used to create enhanced images optimized for nodule visualization (Figure 2A). Common bean root images in both enhanced and RGB format were manually annotated using MakeSenseAI (Skalski, 2019) to generate the training datasets for an object detection model using YOLOv11 (Jocher and Qiu, 2024) (Figure 2B). RGB and enhanced models were trained and evaluated in parallel using identical pipelines (Figure 2A). An independent validation set comprising images of common bean and *Medicago* roots was used to assess model performance.

**Figure 2.**
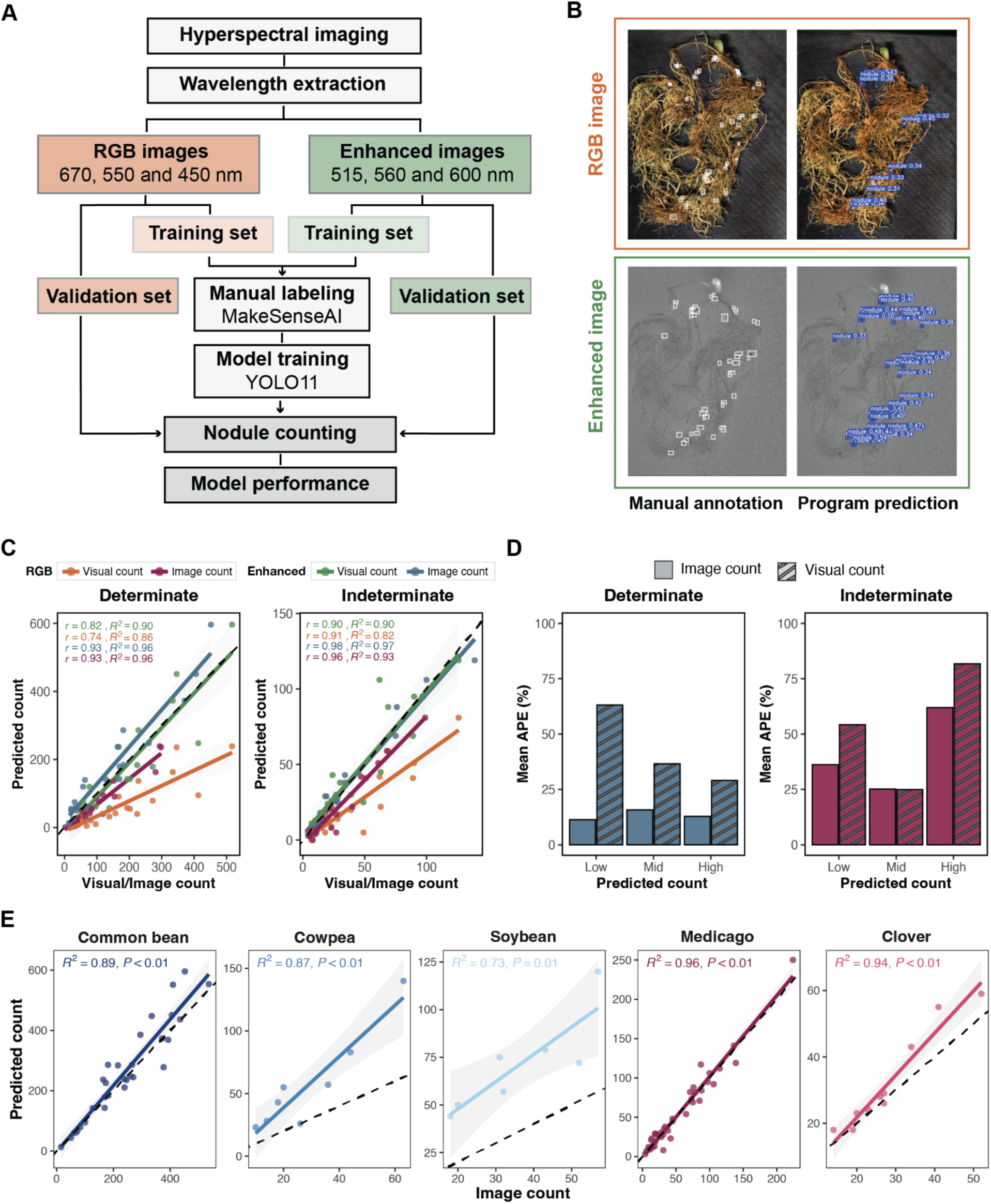
Training a machine learning model to accurately detect nodules using enhanced hyperspectral images and RGB images. (A) Flowchart outlining the steps used in model training and validation. (B) Representative images of manually annotated nodules and model-predicted nodules on enhanced and RGB images of common bean roots. (C) Correlation between model predictions and visual and image counts using enhanced and RGB images (color-coded as indicated in the legend). Pearson correlation coefficients (r) and coefficients of determination (R²) for each group are shown within the plot. (D) Mean absolute percentage error (APE) in detecting determinate and indeterminate nodules using enhanced images. Results are grouped into low, mid and high bins based on predicted count distribution. (APE = abs(Visual/Image count - program count)/Visual/Image count*100). (E) Model performance across different legume species.

This model, trained on common beans (determinate nodules), exhibited poor generalization when applied to *Medicago* (indeterminate nodules), frequently overcounting due to branched indeterminate nodules being incorrectly identified as multiple individual nodules (Figure S3). To resolve this, we trained separate object detection models for each nodule morphotype: a determinate model using common bean root images, and an indeterminate model using *Medicago* root images, both following the same training pipeline (see Table S2 for dataset details). Performance of each model was evaluated by comparing predicted nodule counts (generated by the model) to image counts (manually annotated on the same image) and visual counts (directly observed from the root system by the naked eye). In both models, predicted counts were strongly correlated with manually annotated image counts (R² > 0.93; Figure 2C), confirming that morphotype-specific training substantially enhances detection accuracy.

### Enhanced hyperspectral model outperforms the RGB-based model in detecting nodules

Predicted counts were compared to image counts to assess alignment with manual annotations on the same image, and to visual counts to account for nodules potentially missed due to imaging angle, root density, or other obstructions. For enhanced images, image counts included only active nodules identified by spectral signatures, whereas visual counts included all visible nodules regardless of activity.

Overall, nodule detection trends for both nodule types showed strong positive correlations with both images and visual counts (coefficient of determination (R^2^) > 0.82 and Pearson correlation coefficient (r) > 0.74; Figure 2C). The predicted counts aligned more closely with image counts than visual counts, particularly for indeterminate nodules. The discrepancy between predicted and visual counts was more pronounced in the determinate model, where enhanced images yielded r = 0.93 with image counts and r = 0.82 with visual counts. For both morphotypes, models trained on enhanced images consistently outperformed those using RGB images, likely due to reduced background noise. However, in both models, enhanced images consistently detected more nodules than RGB images, evident in both the quantitative counts (Figure 2C) and during visual inspection (Figure 2B). In the indeterminate model, predictions from both image types aligned more closely with manual counts than in the determinate model, though RGB-based predictions still underestimated nodule numbers (Figure 2C).

Detection accuracy from enhanced images was assessed using mean Absolute Percentage Error (APE) against both image and visual counts (Figure 2D). Overall, error rates were lower for determinate than indeterminate nodule images, and increased when compared to visual counts. Relative to image counts, APE for determinate nodules remained low (12–15%) across all count bins. However, when assessed against visual counts, APE exceeded 50% at low nodule numbers, decreasing with higher counts (Figure 2D). The mean APE relative to image counts was higher for the indeterminate model, ranging from 25% to 60%, and increased with nodule abundance (Figure 2D). As with determinate nodules, APE values were further elevated when predictions were benchmarked against visual counts (Figure 2D).

### Morphotype-specific models enable cross-species nodule detection

To evaluate cross-species applicability, the determinate and indeterminate nodule detection models trained on common bean and *Medicago*, respectively, were applied to soybean, cowpea, and red clover. The determinate model, used for cowpea and soybean, performed well on cowpea, with predicted counts closely tracking image-based counts (R² = 0.87), though overestimation was more pronounced in roots with higher nodule counts (Figure 2E). In contrast, performance on soybean was weaker (R² = 0.73), with consistent overprediction across all count levels, suggesting morphological divergence between the species was not captured by the training data. The indeterminate model applied to red clover showed a strong correlation with image-based counts (R² = 0.94). The model produced more accurate predictions on images with fewer nodules, while a slight overestimation was observed in images with higher nodule counts (Figure 2E). Both models performed optimally on the species used for training. Although model performance varies by species, the two trained models provide a scalable framework for nodule detection across legumes with similar nodule morphotypes.

### Case Study: Nodule detection model links symbiosis in response to environmental cues

To demonstrate the practical and biological relevance of the nodule detection model, we applied it in a controlled experiment simulating real-world environmental responses. Nodulation is closely linked to the plant’s overall nitrogen status, and legumes adjust their symbiotic interactions in response to environmental conditions. Physiologically, it is well known that both soil nitrate levels and atmospheric CO₂ concentrations influence the symbiotic process: high nitrate levels inhibit nodulation (van Noorden et al., 2016), whereas elevated CO₂ promotes it (Libault, 2014; Rogers et al., 2009). Both factors play a role in optimizing the plant’s resource allocation and nutrient use efficiency.

We grew common bean and *Medicago truncatula* under a factorial design incorporating combinations of low/high nitrogen conditions, and ambient/elevated CO₂ levels (Figure 3A, details in Table S3). Root systems were imaged using hyperspectral imaging, and nodule counts were obtained using our nodule detection model. In parallel, visual counts were recorded from the roots to serve as a reference to evaluate model performance. Overall, the nodule detection model produced predicted nodule counts comparable to visual nodule counts, effectively capturing the dynamics of nodulation in response to varying nitrogen and carbon availability in the growth environment (Figure 3B and C). Specifically, both the determinate and indeterminate detection models successfully captured the upregulation of nodulation under elevated CO₂ conditions (Figure 3B). The effect of nitrogen level on downregulation of nodulation was more pronounced in the determinate model, whereas it was present but less apparent in the indeterminate model (Figure 3C).

**Figure 3.**
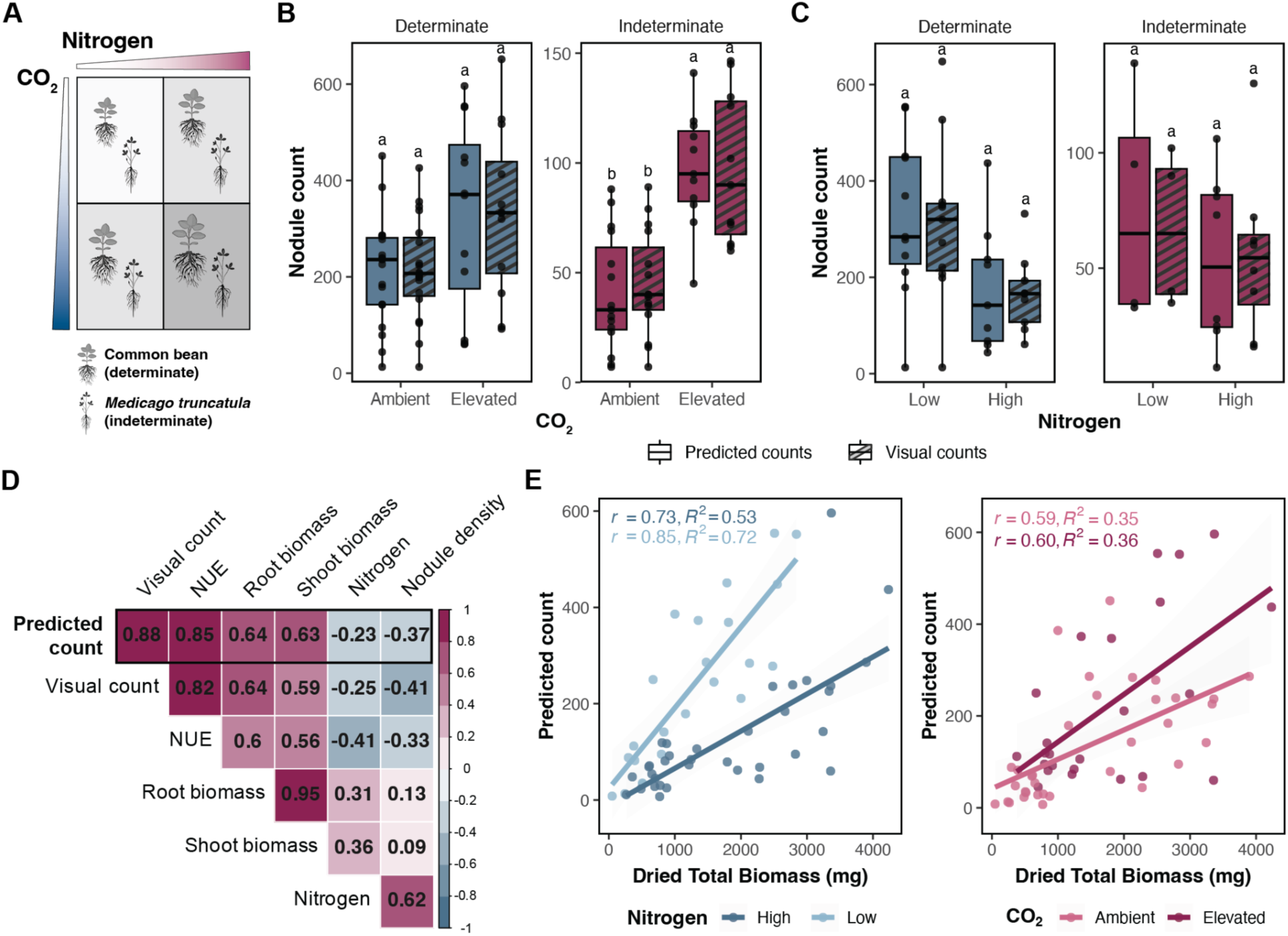
Models trained on hyperspectral-enhanced images accurately capture variation in nodulation patterns under different environmental conditions. (A) Experimental design showing nitrogen and CO₂ treatments applied to bean and Medicago plants. (B) Quantification of nodules under CO_2_ treatments. (C) Quantification of nodules under nitrogen treatments (low = 5 mM NO_3_^-^ and high = 20 mM NO_3_^-^). (D) Correlation of predicted counts with physiological traits. (E) Relationship between predicted nodule counts and total biomass across nitrogen and CO_2_ treatments. Statistical significance in (B) and (C) was assessed using two-way ANOVA followed by Tukey’s post hoc test.

Since our detection model quantifies actively nitrogen-fixing nodules, we further examined the relationship between nodule counts and physiological measurements to assess the contribution of symbiotically assimilated nitrogen to overall plant growth. Pearson correlation analyses were performed between both predicted and visual nodule counts and various physiological traits (Figure 3D). First, we observed a strong correlation between predicted and visual nodule counts, indicating successful detection by our model (Figure 3D). Furthermore, nodule counts showed positive correlations with shoot biomass, root biomass, and NUE, suggesting that greater nodulation is associated with enhanced plant growth and nutrient acquisition (Figure 3D). Nodule counts showed a negative correlation with nitrogen treatment level, providing further evidence of the inhibitory effect of nitrate on nodulation (Figure 3D).

Next, we assessed whether the nodule detection model could capture environmentally driven patterns in nodulation linked to biomass. Under low nitrogen, predicted nodule counts were more strongly correlated with shoot biomass, suggesting greater reliance on symbiosis when nitrogen is limited (Figure 3E). In contrast, elevated CO₂ strengthened this correlation, indicating enhanced benefits from nitrogen fixation (Figure 3E). Together, these results demonstrate that our nodule detection model is well suited for application in research settings, providing a high-throughput, rapid and reliable alternative to the laborious manual counting while also serving as a proxy indicator of a plant’s BNF activity.

## Discussion

Understanding the symbiotic relationship is crucial for future crop improvement (Peoples et al., 1995). Nodule quantification is often used as a rapid reference for estimating BNF activity. However, acquiring such data typically relies on manual data collection by researchers, a process that is time-consuming, labor-intensive, and prone to human error. Here, we present an image-based nodule detection model as an alternative approach for efficient and standardized nodule quantification.

### Spectral wavelength act as standard of nitrogen-fixing nodule detection

Given the importance of nodule quantification in assessing symbiotic performance, researchers have made multiple attempts to standardize the data collection process, with ongoing efforts aimed at improving its efficiency and scalability. Most approaches begin with image-based detection, leveraging nodules’ visual features to identify and quantify nodules in a non-destructive manner. However, these approaches typically rely on semi-automated detection models trained solely on RGB images, with nodule identification still dependent on manual preprocessing by human annotators (Chung et al., 2020; Lira and Smith, 2000; Vikman and Vessey, 1993). Because nodules must be manually marked prior to model training or counting, these methods still require a consistent visual standard for defining what constitutes a nodule. To identify nodules, researchers often rely on visual cues such as physical irregularities or “bumpy” characteristics along the root axis (Chung et al., 2020), or look for circular structures in the images (Hensley et al., 2021). However, such standards are inherently subjective and may limit consistency across different species, particularly when nodules vary in size, shape, and contrast against the root background, or when root orientation differs across images. Moreover, identification based purely on morphological characteristics does not capture the functional status of nodules, and therefore fails to quantify nodulation in terms of symbiotic efficiency (Remmler et al., 2014). To examine the level of nitrogen fixation, destructive chemical assays are required, typically involving nodule dissection, biochemical and isotopic analyses to quantify hemoglobin content (Wilson and Reisenauer, 1963), nitrogenase activity and nitrogen accumulation (Hardy et al., 1968; Vance et al., 1979). However, such methods remain time-consuming, low-throughput, and indirect, as they rely on measurements that do not capture dynamic changes in fixation activity.

In our approach, by utilizing three targeted spectral wavelengths (515 nm, 560 nm and 600 nm), we were able to clearly distinguish nodule tissue from root structures in the hyperspectral enhanced images. We demonstrate that the selected wavelengths are likely indicative of leghaemoglobin content, a key indicator of nitrogen fixation activity (Navascués et al., 2012). By using leghaemoglobin as an anchor for nodule identification, this approach provides a more standardized way of distinguishing nodules from root tissue and offers a more biologically relevant assessment of nitrogen fixation in a non-destructive manner. Furthermore, closer examination of the enhanced images revealed that nitrogen-fixing nodules exhibited varying levels of spectral enhancement, which may correspond to differences in nitrogen-fixing efficiency. Thus, this imaging approach not only accurately distinguishes nodules from root tissue but also holds a promising future for assessing nitrogen fixation activity and capturing developmental dynamics at the level of individual nodules.

### Nodule detection pipeline allows time-efficient high-throughput nodule quantification

Past research has explored various approaches for nodule identification, including manual annotation of images and direct isolation of nodules for downstream counting analysis (Chang et al., 2024; Hensley et al., 2021; Woo et al., 2023). All of these methods require substantial time and labor for sample preprocessing. This often involves carefully spreading out the root systems to avoid overlapping structures and ensuring full exposure of all nodules, which can be particularly challenging under high nodule density or complex root architecture. In contrast, because our model captures nodule-specific spectral wavelengths that remained detectable even under partial root obstruction, the sample preprocessing is significantly simplified, requiring only that roots are reasonably spread out as long as a compatible soil mix is used.

Following the sample preprocessing, many previous studies have demonstrated successful nodule quantification; however, few have compared their program outputs to actual visual counts or evaluated their relationship with physiological traits. Two studies (Lira and Smith, 2000; Vikman and Vessey, 1993) that reported the accuracy of their image-based nodule detection models showed correlation coefficients (r) ranging from 0.85 to 0.89, which is comparable to our model’s performance (r ranging from 0.89 to 0.96), with our model having a simpler sample preprocessing pipeline. Previous automated nodule counting efforts have largely focused on legumes with determinate nodules, with limited adaptation for indeterminate types likely due to their complex, branched morphology. Our morphotype-specific model enables accurate automated quantification of indeterminate nodules (r ≥ 0.90), reporting the first successful implementation and offering scalability across diverse legume species.

While previous studies have explored the use of HSI to segment belowground structures such as roots and nodule-like features under varying conditions (Chang et al., 2024), few have focused specifically on functional nodule identification or quantitative phenotyping using advanced object detection models. Our work demonstrates that integrating HSI with cutting-edge detection algorithms like YOLOv11 offers a scalable, high-throughput, and functionally informative approach to legume nodule phenotyping. The automated nodule counting program developed using YOLOv11 is user-friendly and computationally efficient, requiring only ∼20 milliseconds per image on an NVIDIA A100 GPU <u>(Figure S4</u>). This rapid processing speed supports high-throughput phenotypic analysis with large datasets.

### Model limitations and future directions

Several practices should be considered to ensure optimal detection performance. First, removing soil particles will improve nodule prediction accuracy, especially when using substrates like BM6 potting mix, which will obscure nodule visibility if densely covering the plant tissue, or turface, which shares a similar spectral wavelength as nitrogen-fixing nodules (Figure 1G). Residual fine white silica sand on nodules has minimal impact. Second, the timing of imaging post-harvest is important. Roots should be imaged after cleaning with minimal water; delays longer than 24 hours may lead to drying of nodules and degradation of leghaemoglobin content, which reduces image enhancement quality. Root tissue can be frozen and then thawed before imaging without a negative impact. Third, a high-resolution hyperspectral camera (with at least 900 pixel resolution) is needed for detailed detection of nodules on the images. For researchers without access to a hyperspectral imaging system, an alternative approach is to use narrow-band filters centered at 515 nm, 560 nm, and 600 nm to capture nodule-specific reflectance. By capturing individual images at each of these wavelengths and subsequently combining them, an enhanced composite image can be generated that highlights nodules with improved contrast relative to root tissue.

Based on the comparison between predicted counts and visual counts, our nodule detection model effectively captures the overall trend in nodule number per image. However, variations in nodule size across different species can lead to deviations in the predicted count accuracy (Figure 2E). Tightly clustered nodules often lead to inaccurate counts, as their combined spectral signal can be overly intense, obscuring the boundaries between individual nodules and hiding the possible small nodules behind, resulting in a single, merged detection of a large nitrogen-fixing area. From the case study, the inhibition of nodulation in plants forming indeterminate nodules is less apparent (Figure 3C). Since indeterminate nodules maintain active meristematic tissue (Xiao et al., 2014), the growth and expansion of existing nodules may be favored over the initiation of new ones, leading to smaller observable changes in nodule count. With these limitations in mind, further development of the nodule quantification pipeline can help improve its accuracy and applicability. To address challenges of morphological differences among nodules across species, expanding the training dataset to include a wider range of root images with diverse nodule traits can further enhance the model’s generalizability. An alternative strategy could involve quantifying the total nitrogen-fixing area rather than discrete nodule counts for tightly clustered nodules and quantifying physiological response on indeterminate nodules.

## Acknowledgements

We thank Michelle McReynolds, Ely Lubash and Drew Kim for their assistance with experimental work and Seldon Kwafo for designing and constructing the rhizobox system. We also thank Ana Saballos, Jinwook Kim, Laurie Leonelli, and Katy Heath for providing seeds and rhizobial strains for experiments.

