## Supplementary figures for "Hyperspectral Imaging to Quantify Nodules and Detect Biological Nitrogen Fixation in Legumes"

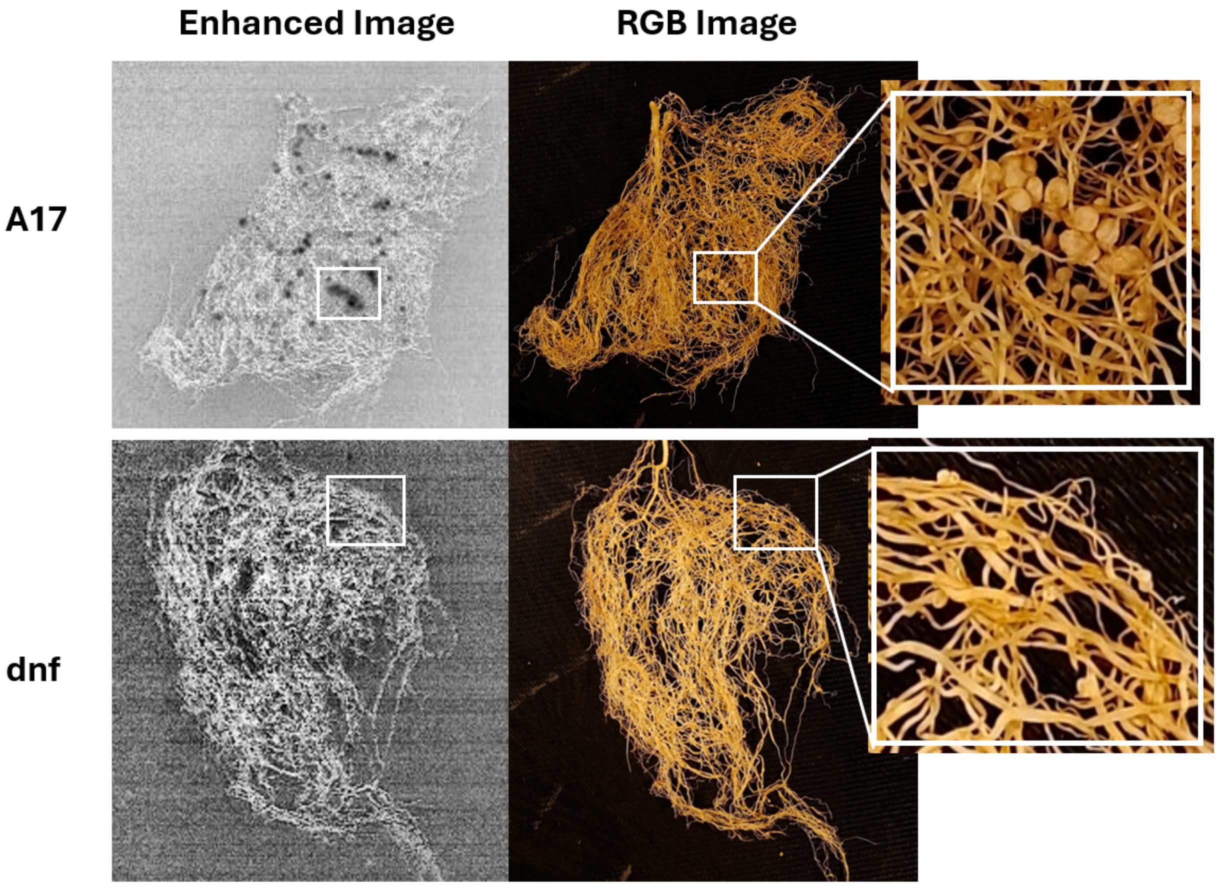


**Figure S1.** Enhanced hyperspectral images of *Medicago truncatula* ecotype A17 and the *dnf* mutant (*does not fix*). A17 represents the wild-type, while *dnf* is an A17-background mutant defective in bacteroid differentiation and lacking leghaemoglobin.

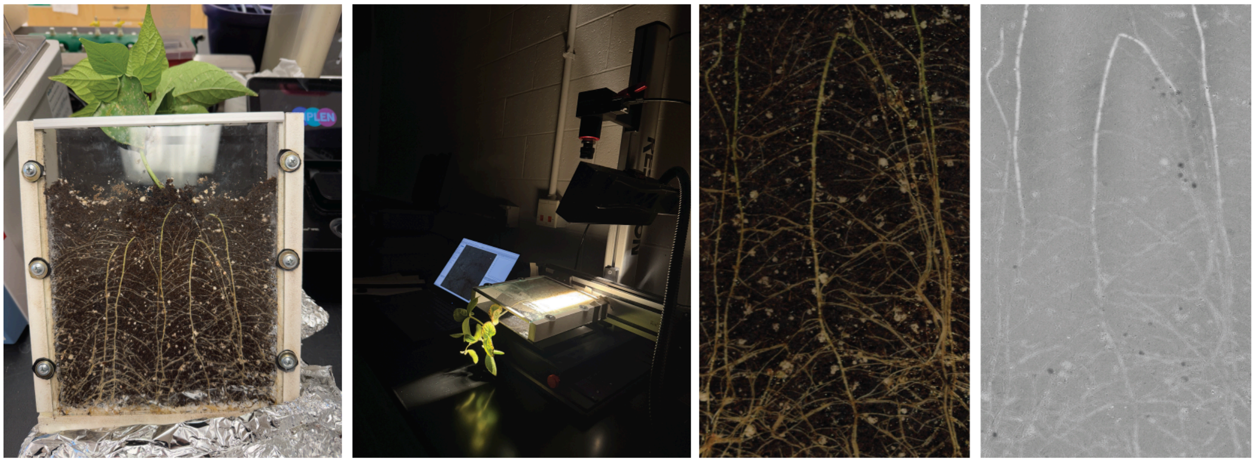


**Figure S2** Rhizobox used to grow plants for root and nodule visualization, imaging setup and extracted RGB and enhanced images.

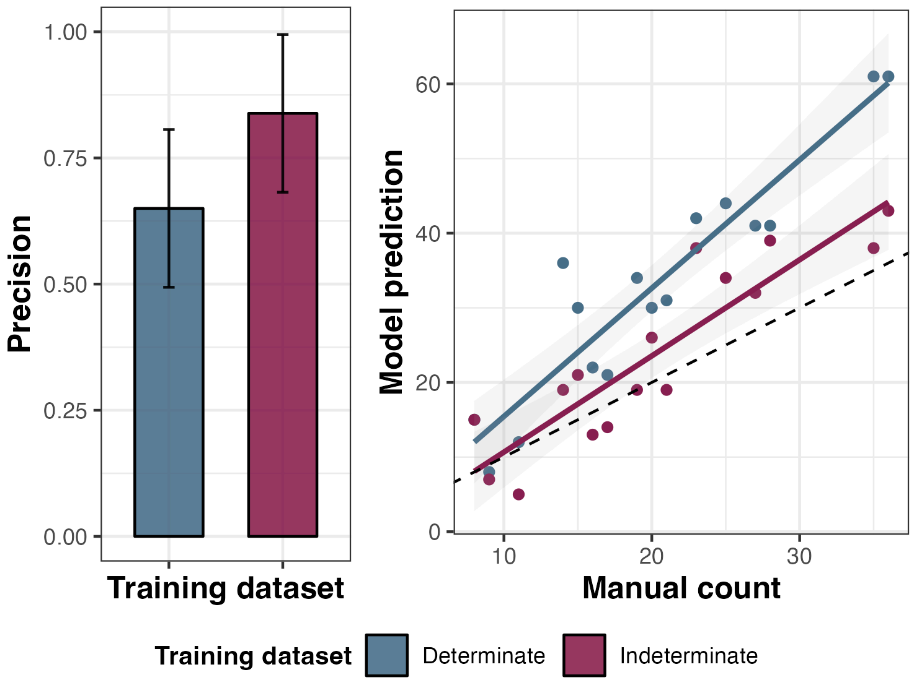


**Figure S3** Validation of indeterminate nodules images of *Medicago truncatula* (n=16) using models trained on determinate nodule images from *Phaseolus vulgaris* and indeterminate nodules images of *M. truncatula*. Precision = min(Manual count/Model prediction,1)

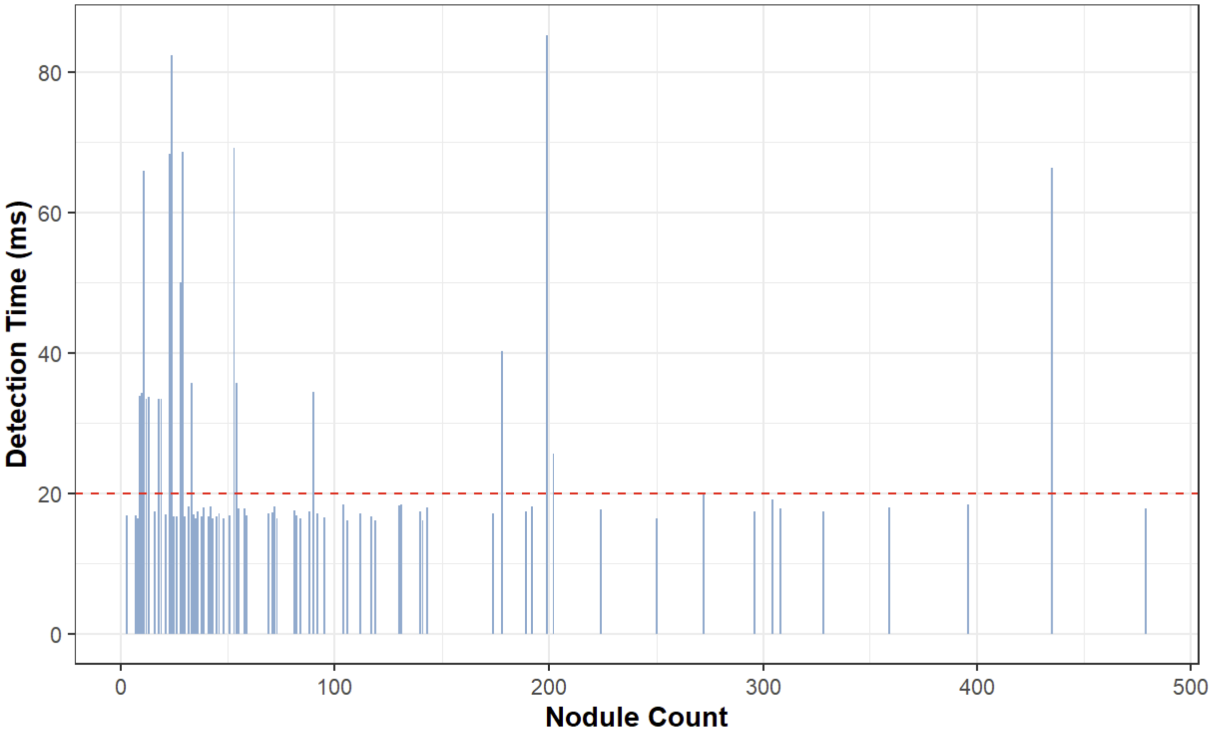


**Figure S4** Nodule detection time required by the counting model. Red dashed line: average of detection time.
